# Sequential combinations of chemotherapeutic agents with BH3 mimetics to treat rhabdomyosarcoma and avoid resistance

**DOI:** 10.1101/2020.01.24.918532

**Authors:** Clara Alcon, Albert Manzano-Muñoz, Estela Prada, Jaume Mora, Aroa Soriano, Gabriela Guillén, Soledad Gallego, Josep Roma, Josep Samitier, Alberto Villanueva, Joan Montero

## Abstract

Rhabdomyosarcoma (RMS) is the most common soft tissue sarcoma in childhood and adolescence. Refractory/relapsed RMS patients present a bad prognosis that combined with the lack of specific biomarkers difficult the development of new therapies. We here utilize dynamic BH3 Profiling (DBP), a functional predictive biomarker that measures net changes in mitochondrial apoptotic signaling, to identify anti-apoptotic adaptations upon treatment. We use this information to guide the use of BH3 mimetics to specifically inhibit BCL-2 pro-survival proteins, defeat resistance and avoid relapse to therapy. Indeed, we found that BH3 mimetics that selectively target BCL-xL and MCL-1 synergistically enhance the effect of the clinically used chemotherapeutic agents vincristine and doxorubicin in RMS cells. We validated this strategy *in vivo* using a RMS patient-derived xenograft (PDX) model and observed a reduction on tumor growth with a tendency to its stabilization with the sequential combination of vincristine and the MCL-1 inhibitor S63845. Finally, we identified the molecular mechanism by which RMS cells acquire resistance to vincristine: through the anti-apoptotic protein MCL-1 for which we observed an enhanced binding between MCL-1 and BID after drug exposure, which is suppressed by the sequential addition of S63845. In conclusion, our findings validate the use of DBP as a functional assay to predict treatment effectiveness in RMS and provide a rationale for BH3 mimetic combination with chemotherapeutic agents to avoid tumor resistance, improve treatment efficiency and decrease undesired secondary effects.

## Introduction

Rhabdomyosarcoma (RMS) is a highly malignant cancer that, despite being relatively rare, is the most frequent soft-tissue sarcoma in children, accounting for 5% of all pediatric tumors^1^. RMS tumors are highly aggressive and typically develop from skeletal muscle cells arising in a variety of anatomic sites in the body^2,3^. There is a slightly higher prevalence of this disease in males than in females, and it is often associated with genetic disorders such as Li-Fraumeni familiar cancer syndrome and neurofibromatosis type 1^2^. Based on histologic criteria, RMS tumors are subdivided in two main groups, embryonal (ERMS) and alveolar (ARMS). ERMS account for 60% of all RMS, affecting children under the age of 10, especially around the head and neck regions^2,3^. ARMS represent approximately 20% of all RMS, occurring mostly in adolescents, frequently localized in the limbs^3,4^. The current treatment strategies for RMS include chemotherapy, radiation, and surgery^4^. Despite treatment improvement for patients with low- and intermediate-risk disease, the survival rates for high-risk patients have not advanced in the last decades^4^. Furthermore, the derived toxicity from current treatments and the lack of biomarkers^5^ highlight the need for new therapies to enhance RMS clinical outcomes.

Therapy causes cancer cells’ death mostly by apoptosis, a process controlled by the BCL-2 family of proteins^6^. Its members are classified based on their structure, BCL-2 homology (BH) domains and their function^6,7^. In brief, the anti-apoptotic proteins (BCL-2, BCL-xL MCL-1 and others) have four BH domains (BH1-BH4) and bind to pro-apoptotic proteins. The pro-apoptotic effector proteins BAX and BAK also contain four BH domains and have the capacity to oligomerize and form pores in the mitochondrial outer membrane. Their function is induced by activator proteins possessing a unique BH3 domain, such as BIM, BID, or PUMA. There is a fourth group of BCL-2 family proteins, the so-called sensitizers, also presenting a unique BH3 domain that cannot directly activate effectors but can inhibit anti-apoptotic members. Sensitizers include BAD, HRK, BIK, NOXA and BMF, among others, and exert a pro-apoptotic effect by competing for specific binding to anti-apoptotic BCL-2 family proteins^7^. Overall, these proteins regulate mitochondrial outer membrane permeabilization (MOMP) and the release of cytochrome c (and other proteins) that represents the point of no return for apoptotic cell death. Importantly, MOMP can be prevented by anti-apoptotic proteins through direct binding to BAX and BAK or activator BH3-only proteins^7^.

Evasion of apoptosis is a hallmark of human cancers and it is often explained by anti-apoptotic proteins’ increased expression^8,9^. In fact, high levels of BCL-2 and MCL-1 have been reported in RMS patients as a pro-survival mechanism^10,11^. Therefore, targeting anti-apoptotic proteins represents a promising therapeutic approach to treat high-risk or relapsed RMS patients^9,12^. In this regard, BH3 mimetics, a novel class of therapeutics that mimic the action of sensitizer BH3-only proteins and selectively inhibit anti-apoptotic BCL-2 family members^7^, could be used to overcome apoptotic resistance. There is an increasing interest on BH3 mimetics due to their therapeutic potential alone or in combination with other treatments, but the main question that clinicians must face is: when and how to use BH3 mimetics as anti-cancer therapies in the clinic^7^. On this subject, the functional assay dynamic BH3 profiling (DBP) can determine in less than 24 hours how effective a treatment will be to engage apoptosis^13^. Briefly, this technology uses synthetic BH3 peptides derived from BCL-2 family proteins to measure how close cells are to the apoptotic threshold or how primed cells are for death. DBP has been successfully used to predict, days to weeks in advance, treatment effectiveness in cell lines, murine models and patient samples^13–17^. In addition to overall susceptibility to apoptosis, DBP can identify cancer cells’ selective dependence on anti-apoptotic proteins and guide BH3 mimetic use to overcome therapy-induced resistance^7^.

Several publications by Fulda and colleagues elegantly demonstrate BH3 mimetics’ therapeutic potential to treat RMS ^12,18–20^ although sequential combination of anti-cancer agents with BH3 mimetics has not been fully assessed. Here we report a new strategy that utilizes low-dose combinations of chemotherapeutic agents with BH3 mimetics to increase current treatments’ efficacy while decreasing therapy-induced toxicity^21^ and anti-apoptotic protection.

## Results

### Novel chemotherapy combinations with BH3 mimetics to increase RMS cytotoxicity

Chemotherapeutic agents are commonly used in clinical protocols for RMS treatment^4^. However, they negatively impact patients with short- and long-term therapy toxicities^22^, and often treatment resistance is acquired by cancer cells^23^. Therefore, we focused on reducing chemotherapeutic dosage to decrease therapy-associated undesired effects. First, we used DBP to analyze the increase in priming after incubation with four standard of care RMS chemotherapeutic agents: the microtubule destabilizing agent vincristine, the alkylating molecule cyclophosphamide, the anthracycline antibiotic with antineoplastic activity doxorubicin and the topoisomerase inhibitor etoposide^24^. We performed DBP on three different RMS cell lines to account for the disease heterogeneity: two ARMS cell lines (CW9019 and RH4) and an ERMS cell line (RD). We observed an increase in priming upon treatment (Δ% priming) after a short incubation with vincristine and doxorubicin, but not with cyclophosphamide or etoposide (Figure 1A and 1C). Using Annexin V and propidium iodide (PI) or DAPI staining, we analyzed by flow cytometry cell death after 96 hours of exposure to the same chemotherapeutic agents as a proof of principle to evaluate the correlation between DBP predictions and later cell death. We observed high levels of cell death (between 40% and 80%) after vincristine treatment and even nearly complete elimination of cells with doxorubicin, but no effect with cyclophosphamide or etoposide, confirming DBP predictions (Figure 1B). We observed a similar trend in the other two RMS cell lines, RD and RH4 (Figure 1D). When we statistically compared Δ% priming and % cell death in all three cell lines, we observed a significant correlation (Figure 1E. left). To further determine how good DBP is as a binary predictor for RMS, we performed a receiver operating characteristic (ROC) curve analysis^25^. We observed that the area under the curve (AUC) for our experiments was 0.81 (Figure 1E. right), indicating that DBP presents a good predictive capacity for chemotherapy cytotoxicity in the RMS cell lines tested.

**Figure 1:**
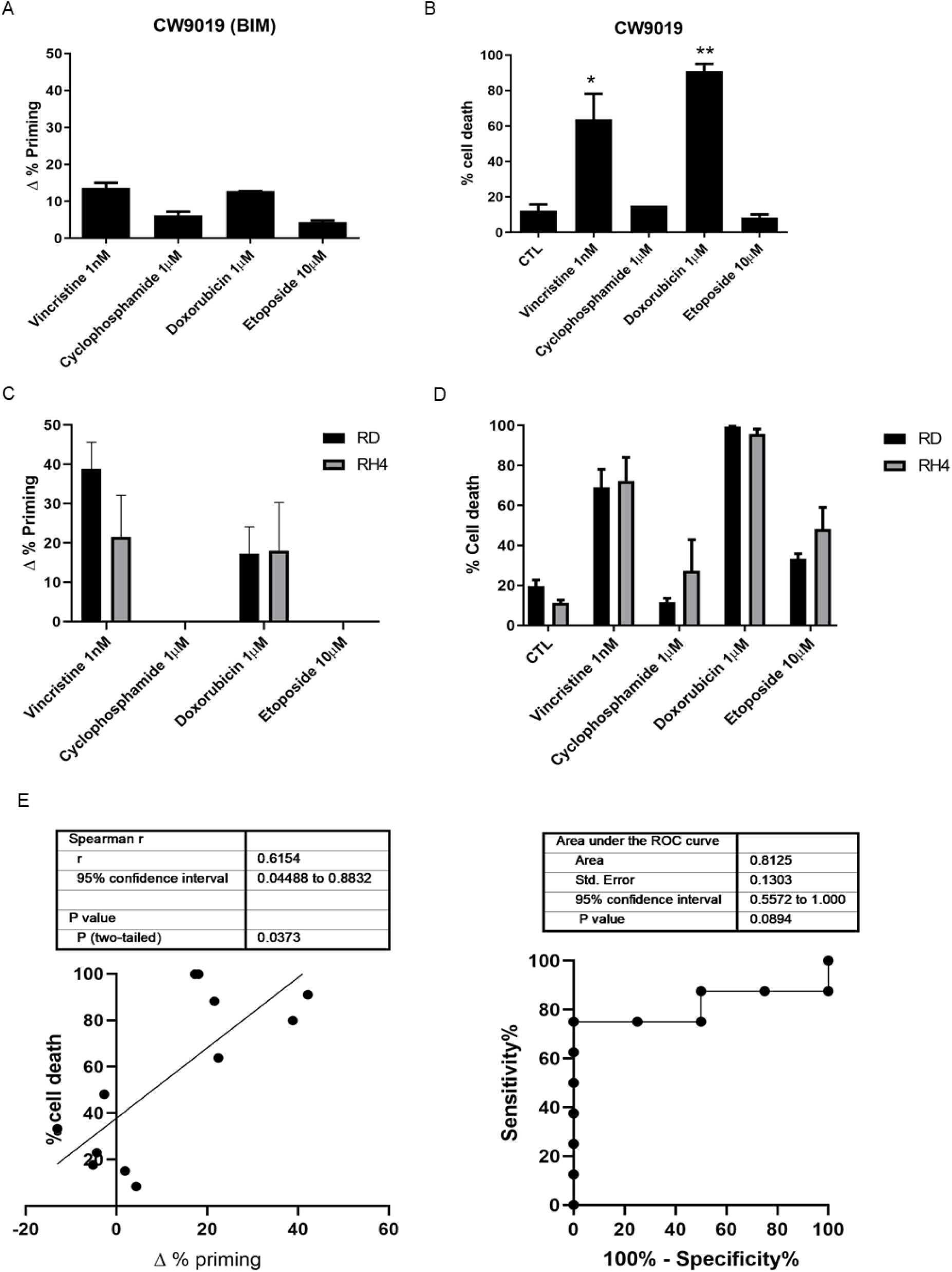
Dynamic BH3 profiling predicts chemotherapy sensitivity in different RMS cell lines. (A) Results from the DBP assay after 36 hours incubation with the treatments in CW9019 cells. Results expressed as Δ% priming represents the increase in priming compared to control cells. (B) Cell death results from Annexin V and propidium iodide/DAPI staining and FACS analysis after 96 hours incubation with the chemotherapeutic agents in CW9019 cells. (C) Results from the DBP assay after 36 hours incubation with the treatments in RD and RH4 cells. Results expressed as Δ% priming represents the increase in priming compared to control cells. (D) Cell death results from Annexin V and propidium iodide/DAPI staining and FACS analysis after 96 hours incubation with the chemotherapeutic agents in RD and RH4 cells. (E) Left plot showing the correlation between Δ% priming at 36 hours and % cell death at 96 hours. Receiver Operating Characteristic curve analysis showed at right. Values indicate mean values ± SEM from at least three independent experiments. ** p<0.01 and * p<0.05.

As mentioned above, one of the hallmarks of cancer is treatment adaptation and resistance to anti-cancer drugs^23^. This resistance can be acquired by different mechanisms such as drug target alterations (mutations), drug export transporters’ gain, increased DNA damage repair, altered proliferation and, as we further investigated, through anti-apoptotic BCL-2 proteins^26^. Using specific synthetic BH3 peptides, that mimic sensitizer BCL-2 family proteins, with DBP we can identify which is the anti-apoptotic protein that cancer cells rely on to acquire resistance to a given treatment^7^. In this regard, we can precisely evaluate the contribution of three main pro-survival BCL-2 family members: BCL-2/BCL-xL dependence with the BAD BH3 peptide, BCL-xL dependence with the HRK BH3 peptide and MCL-1 dependence with the MS1 BH3 peptide^7,27–30^, after treating the cells with a specific therapeutic agent. Using this strategy, we identified that CW9019 cells upon vincristine treatment present an increase in Δ% priming using BAD, HRK and MS1 BH3 peptides (Figure 2A), indicating that cancer cells acquire resistance to this agent mostly through BCL-xL and MCL-1. In consequence, we decided to pharmacologically exploit this anti-apoptotic dependence utilizing two new selective BH3 mimetics: S63845 (MCL-1 inhibitor)^31^ and A-1331852 (BCL-xL inhibitor)^32^ and test their cytotoxic effect in combination with vincristine. We observed that sequentially adding S63845 or A-1331852 after 16 hours of exposure to vincristine significantly increased cell death at 96 hours compared to single agents (Figure 2B). In fact, the combination index (CI) calculations^33^ indicated that S63845 addition to vincristine is synergistic (CI<1) while A-1331852 is additive (CI=1). Obtaining a synergistic combination between two agents is an important goal to decrease treatment toxicity and to avoid undesired side effects associated with high doses of chemotherapy, a constant challenge in pediatric cancer^34^. We repeated these experiments with another RMS standard chemotherapeutic agent, doxorubicin, and we observed an increase in Δ% priming with DBP (Figure 1A) and a high percentage of cell death with Annexin V and DAPI staining (Figure 1B). Like vincristine, we could detect an increase in priming with BAD, HRK and MS1 BH3 peptides in CW9019 cells (Figure 2C) indicating that cancer cells also acquired resistance to doxorubicin treatment through BCL-xL and MCL-1. Doxorubicin is already a potent chemotherapeutic agent as a single agent and exerts an extensive cytotoxicity after 96 hours (Figure 1B), but also causes cardiotoxicity in the clinic^21^. Therefore, we sought to reduce doxorubicin dosing by exploring synergistic sequences with the anti-apoptotic inhibitors A-1331852 and S63845. Hereof, doxorubicin combined with both BH3 mimetics was highly cytotoxic at 96 hours for RMS cells, even when reducing ten-fold its concentration (Figure 2D). Both combinations of doxorubicin with S63845 or A-1331852 were synergistic as we observed a CI < 1. These combinations were effective in RMS cell lines but we did not observe any cytotoxic effect in non-tumoral myoblast cells (Supplementary Figure 1), indicating specific toxicity of these treatments for malignant cells. Additionally, we analyzed several BCL-2 family proteins expression to determine molecular fluctuations after vincristine and doxorubicin treatments. Surprisingly, we found that upon vincristine treatment there were no significant changes in anti-apoptotic proteins MCL-1, BCL-xL or BCL-2 expression, indicating that cancer cells’ adaptation to this treatment relies on different mechanisms (Supplementary Figure 2). On the other hand, doxorubicin treatment lead to a marked decline in MCL-1 and BCL-xL levels, an increase in BCL-2 expression and a decrease in pro-apoptotic proteins BAK and BID (Supplementary Figure 2). From this first set of experiments we conclude that we can increase chemotherapeutic agents’ efficacy by rationally combining them with specific BH3 mimetics. Particularly, sequential vincristine treatment followed by S63845 stands out as the most effective therapy *in vitro* for CW9019 RMS cells (Figure 2B).

**Figure 2:**
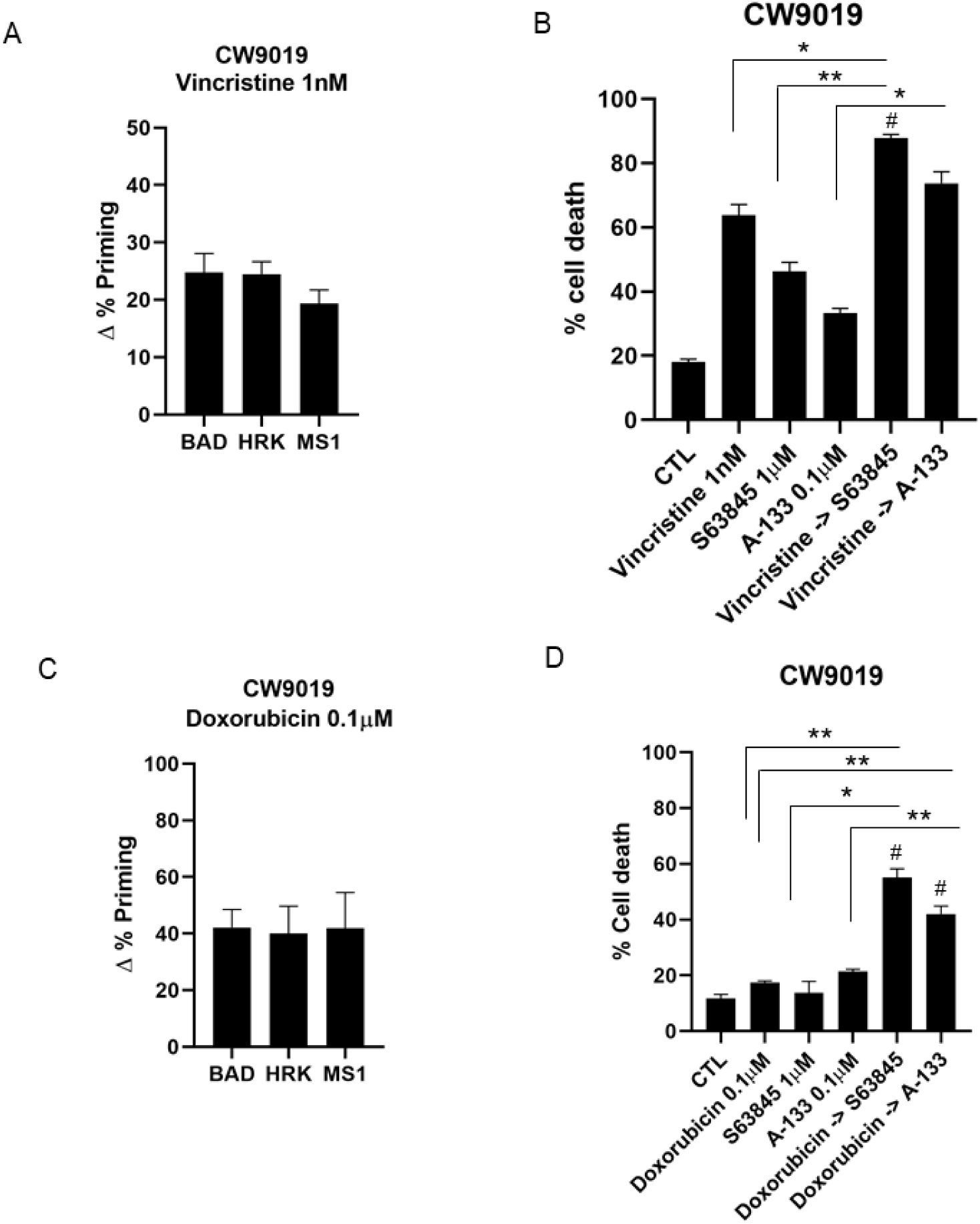
Dynamic BH3 profiling predicts synergistic combinations with BH3 mimetics in CW9019 cell line. (A) Results from the contribution of each anti-apoptotic protein: BCL-2/BCL-xL dependence BAD peptide; BCL-xL dependence HRK peptide; and MCL-1 dependence MS1 peptide in acquiring resistance to vincristine. Results expressed as Δ% priming represents the increase in priming compared to control cells. HRK and MS1 BH3 peptides showed a significant increase, indicating BCL-xL and MCL-1 adaptation respectively (B) Cell death from Annexin V and propidium iodide/DAPI staining and FACS analysis after 96 hours incubation of CW9019 cells with the single agents alone or the combination of vincristine with the corresponding BH3 mimetics S63845 and A-1331852 for 96 hours. (C) Results from the contribution of each anti-apoptotic protein. HRK and MS1 BH3 peptides showed a significant increase in priming, indicating BCL-xL and MCL-1 adaptation respectively (D) Cell death from Annexin V and propidium iodide/DAPI staining and FACS analysis after 96 hours incubation of CW9019 cells with the single agents alone or the combination of doxorubicin with the BH3 mimetics S63845 and A-1331852. Values indicate mean values ± SEM. ** p< 0.01, * p<0.05 compared to single agents and # indicates CI<1. All experiments were performed at least three times.

### Vincristine induces BID sequestering by MCL-1 as a drug-induced resistance mechanism

In order to understand the molecular adaptation by which cells acquire resistance to vincristine and why its combination with S63845 is highly effective *in vitro,* we immunoprecipitated MCL-1 from CW9019 control cell lysates and CW9019 cells treated with vincristine for 36 hours (Figure 3B). MCL-1 can bind to BID and prevent the activation of BAX and BAK, therefore inhibiting cytochrome c release from the mitochondria and apoptosis^35^. We hypothesized that this molecular interaction could explain resistance to vincristine treatment. Thus, we immunoprecipitated MCL-1 and observed a significant protein decrease in the lysate unbound fraction (Figure 3A), but a good detection in the pulled down samples (Figure 3C). When we checked for BID co-immunoprecipitation, we observed that 36 hours after treatment with vincristine, there was a significant increase in MCL-1 and BID binding compared to the control (Figure 3C), explaining apoptosis protection. This increase in binding could be reversed by the addition of S63845 after 16 hours of treatment with vincristine (Figure 3C), recovering the apoptotic function in CW9019 cells. This explains why the sequential combination of vincristine and S63845 is effective against RMS.

**Figure 3:**
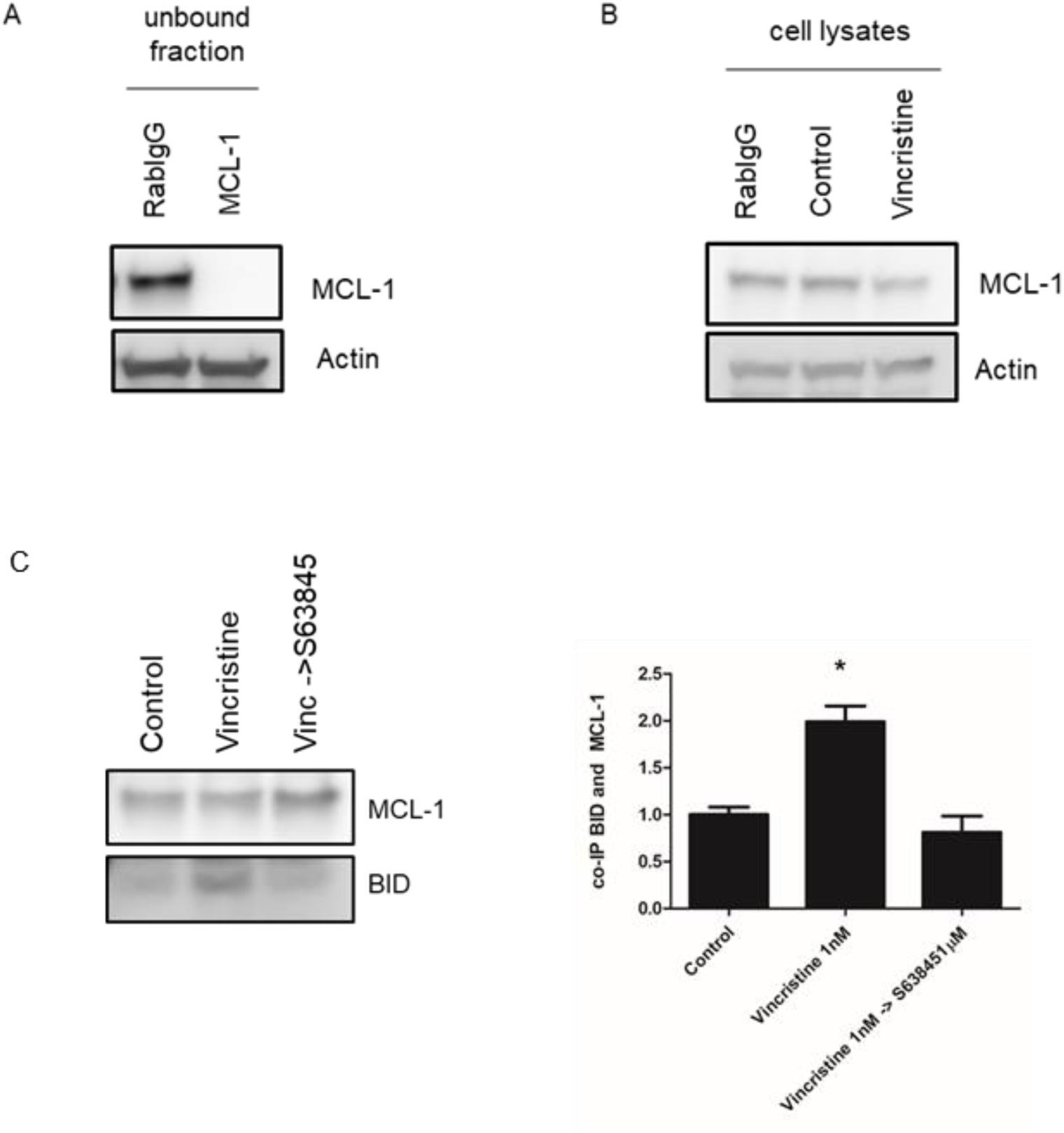
Vincristine induces resistance in RMS cells through BID inhibition by MCL-1. (A) Western blot results of the unbound fraction after MCL-1 immunoprecipitation. High efficiency of MCL-1 immunoprecipitation compared to Rabbit IgG control antibody. (B) Western blot results showing MCL-1 levels in CW9019 cell lysates (incubated with vincristine or DMSO for 36 hours) before performing the immunoprecipitation. (C) Left panel: Western blot results of the co-immunoprecipitation between MCL-1 and BID. Right panel: Quantification of the optical density of each protein and represented as binding ratio between BID and MCL-1. Results showed a significant increase in BID and MCL-1 binding after vincristine treatment, which is restored to control levels after the addition of S63845. Values indicate mean values ± SEM from at least three independent experiments, * p<0.05.

### Effective therapeutic combination *in vivo* of vincristine with the MCL-1 inhibitor S63845

Patient-derived xenografts (PDXs) are advantageous in pre-clinical research as they recapitulate patients’ therapeutic response^22^. After identifying different effective combinations *in vitro*, we analyzed tumors from RMS PDX models. We disaggregated the tumors to obtain a single-cell suspension and perform DBP analyses to evaluate different therapies’ effectiveness and possible anti-apoptotic adaptations. We focused on chemotherapeutic agents, particularly on vincristine as it is utilized in the clinic to treat RMS and we already generated promising preliminary results *in vitro* in combination with S63845 (Figure 1). We analyzed an embryonal rhabdomyosarcoma orthoxenograft generated from a small biopsy from the primary tumor located in the child gluteus of a metastatic child, named RMSX1. We detected an increase in Δ% priming after incubating tumor cells with vincristine (Figure 4A), but not with cyclophosphamide or etoposide (Supplementary Figure 3A) as we previously observed in CW9019, RD and RH4 cell lines (Figure 1). These observations illustrate the great heterogeneity observed in RMS tumors and variable response to chemotherapy in the clinic. RMSX1 treatment with vincristine *in vivo* merely delayed tumor growth after 21 days (Figure 4C). Furthermore, we identified by DBP an anti-apoptotic adaptation mediated by MCL-1 (Figure 4B), that could diminish the efficacy of this chemotherapeutic agent, as previously observed *in vitro* (Figure 2). To confirm these results, we treated RMSX1 mice with the MCL-1 inhibitor S63845 as a single agent, or right after vincristine treatment to overcome apoptotic resistance. Surprisingly, we detected that sequentially combining vincristine and S63845 was significantly more effective than single agents and promoted tumor reduction *in vivo* (Figure 4C and Supplementary Figure 3C). Moreover, we could also observe a significant increase in Δ% priming after incubating tumor cells with targeted agents such as S63845, ABT-199 and SP2509 (Supplementary Figure 3A) and we also identified possible anti-apoptotic adaptations to those treatments by BCL-xL and MCL-1 (Supplementary Figure 3B) that we will further explore. Overall, these results demonstrate that DBP can be used to design more effective therapeutic strategies to overcome anti-apoptotic resistance and stop cancer progression using BH3 mimetics.

**Figure 4:**
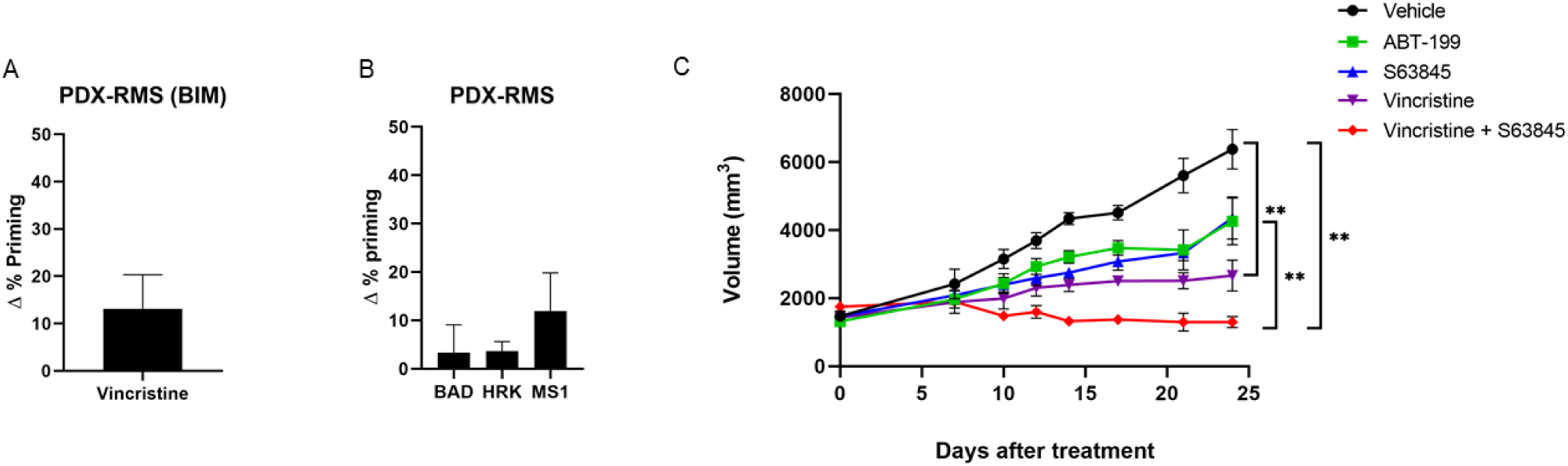
Sequential treatment of vincristine and S63845 stops tumor progression in the PDX model of RMS RMSX1. (A) DBP results of PDX cells from RMS cancer patient showing an increase in Δ% priming after vincristine treatment. Results expressed as Δ% priming represents the increase in priming compared to control cells. n=2 (B) DBP results of PDX cells from RMS cancer patient with the sensitizer peptides. MS1 BH3 peptide showed a significant increase, indicating MCL-1 adaptation. n=2 (C) Tumor growth results after 21 days of treatment with vehicle, vincristine, the BH3 mimetics S63845 and ABT-199 and the combination of vincristine and S63845. Day 0 indicates the day animals received the treatments. All values indicate mean values ± SEM. ** p< 0.01, n=3.

## Discussion

There is an urgent medical need to find more effective and less toxic treatments for RMS patients, since recurrent RMS presents poor prognosis and the overall survival after relapse is very low^28^. There is a growing evidence that the BCL-2 family of proteins (particularly the anti-apoptotic members) may mediate drug resistance in cancer cells promoting patients’ disease progression^7,8^. Therefore, it is key to predict these acquired pro-survival mechanisms and overcome them with anti-apoptotic inhibitors like BH3 mimetics. As mentioned above, dynamic BH3 profiling (DBP), beyond measuring a given treatment effectiveness to engage apoptosis, can also detect anti-apoptotic adaptations derived from therapy that ensure cancer survival^13^ to guide BH3 mimetics use to avoid resistance. Anti-apoptotic inhibitors such as A-1331852 (BCL-xL selective), ABT-199 (BCL-2 selective), S63845 (MCL-1 selective) among others that are now evaluated in clinical trials^7,27^, can be used as single agents or especially in combination with other therapies to enhance cancer cells elimination^7^. Therefore, in this study we investigated BH3 mimetics’ use to boost RMS sensitivity to current chemotherapy.

At present, radiotherapy, surgery and chemotherapy are the standard-of-care for RMS treatment. Regarding the latter, a three-drug combination is currently utilized: vincristine, actinomycin D and cyclophosphamide (VAC). This regimen has become the basis for RMS therapy with the incorporation of other agents such as etoposide, doxorubicin, ifosfamide, cisplatin and others for intermediate risk patients, with scarce clinical outcome improvement^36^. However, secondary effects derived from chemotherapy administration in children are severe and may include infertility, cardiomyopathy or the appearance of secondary neoplasia^36^. One explanation for these unbearable therapy-associated pediatric toxicities rely on differential apoptotic priming between young and adult tissues^21^. Traditional chemotherapy has reached an efficacy plateau in RMS making development of new therapies that increase efficacy while decreasing toxicity a clear unmet need. Thus, we sought to identify possible mechanisms of resistance to chemotherapeutic agents, such as vincristine or doxorubicin that could explain the limited clinical efficacy by analyzing anti-apoptotic changes with DBP (Figure 2). Indeed, we identified anti-apoptotic adaptations to common therapies and tested them in combination with BH3 mimetics to achieve a high cytotoxic effect, around 80%, while decreasing ten-fold their concentration, thus their potential secondary effects. More precisely, we determined novel synergistic combinations of vincristine with the MCL-1 inhibitor S63845, and doxorubicin with the same BH3 mimetic or the BCL-xL inhibitor A-1331852 (Figure 2), the effectiveness of this last combination also observed in osteosarcoma^37^. These three new combinations were synergistic as assessed by CI index (CI<1) and allowed dosing reduction^36^. These results strengthen the importance of BCL-xL and MCL-1 as therapeutic targets in pediatric cancer and specifically in RMS as it has also been recently reported^38^. These treatments have been explored in multiple adult cancers^27,39^, but not in pediatric cancers where current treatments present low effectiveness especially in high risk and relapsed RMS patients^36^. Previous studies in RMS also demonstrated that different BH3 mimetics can potentiate chemotherapeutic treatment effectiveness^18^ when combined with an ATP-competitive mTOR inhibitor^20^ or a histone deacetylase inhibitor^12^ which supports exploring these therapies as new approximations to treat pediatric patients.

As previously mentioned, vincristine is currently used for RMS treatment^36^, but we observed that cells acquire resistance through the anti-apoptotic protein MCL-1. Therefore, we focused our efforts on testing vincristine effectiveness in combination with the MCL-1 inhibitor S63845 *in vivo*. As patient-derived xenografts accurately model patients’ outcome^22^, we used a RMS PDX model to test the sequential combination of low dose vincristine therapy with S63845. First, we confirmed using DBP in *ex vivo* PDX-isolated cancer cells that vincristine resistance was mediated through MCL-1 (Figure 4B), correlating with our previous observations *in vitro* (Figure 2A-B). When a combination of vincristine followed by S63845 was sequentially administered, these PDXs showed a significant reduction on tumor growth with a tendency to its stabilization (Figure 4C-supplementary Figure 3C), in accordance with the high cytotoxicity observed *in vitro* (Figure 2B). To further explain this combination efficacy, we analyzed MCL-1 and NOXA expression but we could not detect significant changes on those proteins (Supplementary Figure 2 and data not shown), pointing to another anti-apoptotic mechanism driving the acquired resistance to vincristine. MCL-1 can exert its anti-apoptotic function by sequestering BID, thus inhibiting BAX and BAK activation and avoiding apoptosis^35^. There are different studies focused on trying to develop molecules that could disrupt the interaction between MCL-1 and BID and therefore restore apoptosis^40,41^. We observed a significant increase in MCL-1 binding to BID by co-immunoprecipitation after 36 hours treatment with vincristine, which was reversed when adding S63845 (Figure 3). This increase in binding between MCL-1 and BID explains CW9019 acquired resistance to vincristine and the high efficiency of sequentially combining this chemotherapeutic agent with S63845 both *in vitro* and *in vivo.* S63845 disrupts the interaction between MCL-1 and BID, by blocking the first, to restore apoptosis in these cells,

In summary, the work that we here present demonstrates DBP’s capacity to predict days in advance the cytotoxic effect of specific treatments in RMS cells. More interestingly, it can identify how RMS cancer cells acquire resistance to therapy and new approaches to overcome it. To our knowledge, this is the first time that multiple effective sequential combinations of chemotherapeutics with BH3 mimetics are reported for RMS in the same study. Indeed, we demonstrated *in vitro* and *in vivo* the synergistic antitumor activity of the MCL-1 inhibitor S63845 when sequentially combined to vincristine. These findings, together with the current efforts to target the anti-apoptotic protein MCL-1, currently explored in clinical trials^42^, manifest the importance of rationally combining anti-cancer agents with BH3 mimetics. These novel therapeutic strategies could improve RMS patients’ treatment in the clinic, also relapsed, when guided by a functional predictive biomarker such as dynamic BH3 profiling.

## Materials and Methods

### Cell lines and treatments

RMS cell lines (CW9019, RD and RH4) were kindly provided by Dr. Oscar Martínez-Tirado and Dr. Cristina Muñoz-Pinedo from the Biomedical Research Institute from Bellvitge (IDIBELL). Cells were tested for mycoplasma and cultured in RPMI 1640 medium (Gibco, 31870) supplemented with 10% heat inactivated fetal bovine serum (Gibco, 10270), 1% of L-Glutamine (Gibco, 25030) and 1% of penicillin and streptomycin (Gibco, 15140) and maintained at 37°C in a humidified atmosphere of 5% CO_2_. Drug treatments were performed directly in the culture media at the doses and time points indicated in every single experiment.

### Dynamic BH3 Profiling

3 × 10^4^ cells/well in a 96-well plate were used for cell lines. 25 μL of BIM BH3 peptide (final concentration of 0.01, 0.03, 0.1, 0.3, 1, 3 and 10 μM), 25 μL of BAD BH3 peptide (final concentration of 10 μM), 25 μL of HRK BH3 peptide (final concentration of 100 μM) and 25 μL of MS1 BH3 peptide ^30^ (final concentration of 10 μM) in MEB (150mM mannitol, 10 mM hepes-KOH pH 7.5, 150 mM KCl, 1 mM EGTA, 1 mM EDTA, 0.1% BSA, 5 mM succinate) with 0.02% digitonin were deposited per well in a 96-well plate (Corning, 3795). Single cell suspensions were stained with the viability marker Zombie Violet (BioLegend 423113) and then washed with PBS and resuspended in MEB in a final volume of 25 μL. Cell suspensions were incubated with the peptides for 1 hour following fixation with formaldehyde and staining with cytochrome c antibody (BioLegend, Alexa Fluor® 647 anti-Cytochrome c - 6H2.B4, 612310). Individual DBP analysis were performed using triplicates for DMSO, alamethecin (Enzo Life Sciences, BML-A150-0005), the different BIM BH3 concentrations used, BAD, HRK and MS1 BH3 peptides. The expressed values stand for the average of three different readings performed with a high-throughput flow cytometry SONY instrument (SONY SA3800). % priming stands for the maximum % cytochrome c released obtained from different BH3 peptide and Δ% priming stands for the maximum difference between treated cells minus non-treated cells.

### Cell death analysis

Cells were stained with fluorescent conjugates of Annexin V (BioLegend, FITC Annexin V, 640906 or Alexa Fluor® 647 Annexin V, 640912) and propidium iodide (PI) (BioVision, 1056) or DAPI (Thermo Fisher, 62248) and analyzed on a flow cytometry Gallios instrument (Beckman Coulter). Viable cells are Annexin V negative and PI or DAPI negative, and cell death is expressed as 100%-viable cells.

### Protein extraction and quantification

Proteins were extracted by lysing the cells during 30 minutes at 4°C using RIPA buffer (150 mM NaCl, 5 mM EDTA, 50 mM Tris-HCl pH=8, 1% Triton X-100, 0.1% SDS, EDTA-free Protease Inhibitor Cocktail) followed by a centrifugation at 16,100 × *g* for 10 minutes. The supernatant was stored at −20°C for protein quantification performed using Pierce ™ BCA Protein Assay Kit (Thermo Fisher, 23227).

### Immunoprecipitation

Cells were lysed in Immunoprecipitation buffer (150 mM NaCl, 10 mM Hepes, 2 mM EDTA, 1% Triton, 1.5 mM MgCl2, 10% glycerol, protease inhibitor from Roche) and centrifuged at 14,000 × *g*, 15 minutes at 4°C. The resulting supernatants were incubated with magnetic beads (Bio-Rad, 161-4021) conjugated to rabbit anti-MCL-1 antibody (5µg, Cell Signaling, CST94296) or Rabbit IgG antibody (5μg, Cell Signaling, CST2729) at 4°C overnight. A fraction of the supernatant (30 μL) were removed and mixed with half volume of 4X SDS-PAGE sample buffer, heated at 96°C for 5 minutes and stored at −80°C as input fractions. After magnetization, a part of the supernatant was mixed with half volume of 4X SDS-PAGE sample buffer, heated at 96°C for 5 minutes and stored at −80°C as unbound fractions. The rest of the supernatant was discarded. The resulting pellet was washed and mixed with 40 µL 4X SDS-PAGE sample buffer and heated 10 minutes at 70°C to allow the dissociation between the purified target proteins and the beads-antibody complex. Finally, sample was magnetized and the supernatant was collected and stored at −80°C as IP fractions for further western blot analysis.

### Immunoblotting

Proteins were separated by SDS-PAGE (Mini-Protean TGX Precast Gel 12%, Bio-Rad, 456-1045) and transferred to PVDF membranes (Amersham Hybond, 10600023). Membranes were blocked with dry milk dissolved in Tris Buffer Saline with 1% Tween 20 (TBST) for 1 hour and probed overnight at 4°C with the primary antibodies of interest directed against: rabbit anti-BCL-2 (Cell Signaling, CST4223), rabbit anti-BCL-xL (Cell Signaling, CST2764), rabbit anti-MCL-1 (Cell Signaling, CST5453), rabbit anti-NOXA (Cell Signaling, CST14766), rabbit anti-BIM (Cell Signaling, CST2933), rabbit anti-Actin (Cell Signaling, CST4970) followed by Anti-rabbit IgG HRP-linked secondary antibody (Cell Signaling, CST7074) in 3% BSA in TBST for 1 hour at room temperature. Immunoblots were developed using Clarity ECL Western substrate (Bio-Rad, 1705060). When necessary, immunoblots were stripped in 0.1 M glycine pH 2,5, 2% SDS for 40 minutes and washed in TBS. Bands were visualized with LAS4000 imager and ImageJ was then used to measure the integrated optical density of bands.

### Animals and human tissue

Six-week-old male athymic nu/nu mice (Envigo) weighing 18–22 g were used in this study. Animals were housed in a sterile environment, in cages with autoclaved bedding, food, and water. The mice were maintained on a daily 12 hours light, 12 hours dark cycle. The patient gave written consent to participate in the study. The Institutional Ethics Committees approved the study protocol, and the animal experimental design was approved by the IDIBELL animal facility committee (AAALAC Unit1155). All experiments were performed in accordance with the guideline for Ethical Conduct in the Care and Use of Animals as stated in The International Guiding Principles for Biomedical Research Involving Animals, developed by the Council for International Organizations of Medical Sciences.

### Development of rhabdomyosarcoma orthoxenograft mouse model

An embryonal rhabdomyosarcoma (ERMS) orthoxenograft was generated from a small biopsy of a metastatic case taken at diagnostic from the primary tumor located in the child gluteus of a metastatic child. The primary tumor did not receive radiotherapy or chemotherapy prior to surgery. Under isoflurane anesthesia, a subcutaneous pocket was made with surgical scissors. Then, a small incision was made in the muscle and the tumor was fixed with synthetic monofilament, non-absorbable polypropylene suture (Prolene 7.0) to the muscle of the upper thigh (orthotopic implantation). After implantation, tumor formation was checked weekly by palpation. Orthotopic tumor (named RMSX1) became apparent 1–3 months after engraftment. Once orthotopic tumors had reached a volume of around 1,500 mm3, mice were sacrificed and tumors were passed to another three animal in order to obtain a sufficient quantity of tumor material. After each passage tumors were frozen, paraffin-embedded, and cryopreserved in (10% DMSO + 90% Fetal Bovine Serum (FBS) (non-inactivated) to provide a source of viable tissue for future experiments.

### Drug treatment in embryonal rhabdomyosarcoma RMSX1 orthoxenograft tumor model

The orthoxenograft procedure was approved by the campus Animal Ethics Committee and complied with AAALAC (Association for Assessment and Accreditation of Laboratory Animal Care International) procedures. A mouse harboring RMSX1 tumor orthotopically growing (at passage#2) was sacrificed, tumors were harvested and cut into small fragments 4× 4 mm^3^, and the tumor fragments were grafted in 20 young mice. When tumors reached a homogeneous size (1,200 to 1,500 mm^3^) mice were randomly allocated into the different treatment groups (n=4/group): i) Placebo; ii) ABT-199 (100 mg/kg); iii) vincristine (1 mg/kg); iv) S63845 (20 mg/kg); and v) combined vincristine (1 mg/kg) plus S63845 (20 mg/kg). Vincristine was intravenous administrated by tail vein injection (i.v) once per week for 3 consecutive weeks (days 0, 7, and 14). ABT-199 was daily administered (q.d) by oral gavage (p.o) for 21 days and S63845 was i.v administered three consecutive days per week for 2 weeks. All the animals/groups were sacrificed at day 21. To minimize in combined treatments the risk to develop drug induced toxicity drugs were administered spaced in time. Vincristine was administered first and S63845 2 hours later. Vincristine from Eli Lilly (1 mg/ml) was purchased at the hospital pharmacy of Catalan Institute of Oncology (ICO) and diluted in saline before use. ABT-199 and S63845 were purchased at Selleckchem. ABT-199 was diluted in 10% Ethanol/30% PEG 400/60% Phosal 50 PG (v/v/v) while S63845 was diluted in 10% DMSO/40% PEG 300/5% Tween 80/Saline. After treatment initiation, tumors were measured using a caliper every 2–3 days and tumor volume was calculated using the formula v = (w2 l/2), where l is the longest diameter and w the width. At sacrifice, tumor was dissected out and weighed. Representative fragments were either frozen in nitrogen or fixed and then processed for paraffin embedding.

### PDX cell isolation

Primary tumors from PDX animals were exposed to an enzymatic digestion after, mechanical disaggregation, in 2.5 mL of DMEM media with 125 units of DNAse I (Sigma-Aldrich, DN25), 100 units of Hyaluronidase (Sigma-Aldrich, H3506) and 300 units of collagenase IV (Gibco, 17104–019). The tissue suspension was processed using gentleMACS Dissociator (Miltenyl Biotec) using the hTUMOR 1 program. The suspension was incubated at 37°C for 30 minutes in constant agitation. After the program hTUMOR 1 was ran again and repeated the 30 minutes incubation. We filtered the suspension 70 micron filter into a 50 mL conical and cells were spinned down 500 × *g* for 5 minutes. To lyse the residual red blood cells, 100 µL of ice cold water was added for 15 s and then diluted to 50 mL with PBS, then spin cells down again. Cells were finally resuspended in RPMI media, counted by trypan blue exclusion and plated in a 12-well plate, 3 × 10^4^ cells/well and treated with DMSO or vincristine 1 nM. After a 16 hours incubation at 37°C in a humidified atmosphere of 5% CO_2_. Dynamic BH3 profiling analyses were then performed.

### Statistical analysis

Statistical analysis P values 0.05 were considered as statistically significant and results were analyzed using Student’s t-tail test. SEM stands for Standard Error of the Mean. For ROC curve analysis cell lines were considered responsive to treatment when Δ% cell death > 20 %. Drug synergies were established based on the Bliss Independent model as previously described ^33^. Combinatorial index (CI) was calculated CI= ((D_A_+D_B_)-(D_A_*D_B_))/D_AB_, where D represents cell death of compound A or B or the combination of both. Only the combination of drugs with a CI<1 were considered synergies. GraphPad Prism8 was used to generate the graphs and to perform the statistical analysis.

## Acknowledgments

We would like to thank to Dr. Martinez-Tirado and Dr. Muñoz-Pinedo for providing the cell lines used in this study. We would also like to thank the Cytometry Facility from the University of Barcelona for assistance with flow cytometry experiments. This work was supported by the CELLEX foundation and the FERO foundation. J.Montero acknowledges the Ramon y Cajal Programme, Ministerio de Economia y Competitividad (RYC-2015-18357).

## Author contributions

C. Alcon performed all *in vitro* experiments, performed the PDX tumors disaggregation and their DBP analyses. A. Villanueva and A. Soriano performed and analyzed the *in vivo* experiments. G. Guillén, A. Soriano, S. Gallego and J. Roma provided the primary tumors to generate the RMS PDX model for *in vivo* experiments. E. Prada and J. Mora provided some reagents used in the article and expertise. J. Samitier and J. Montero supervised the work. C. Alcon and J. Montero wrote the manuscript.

