## Supplementary figures for "Sequential combinations of chemotherapeutic agents with BH3 mimetics to treat rhabdomyosarcoma and avoid resistance"

### Figure 1

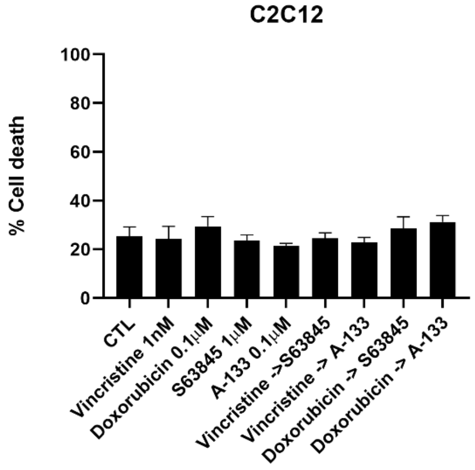


***Supplementary Figure 1: Chemotherapeutics combined with BH3 mimetics have no effect in C2C12 cell line.*** *Cell death from Annexin V and propidium iodide/DAPI staining and FACS analysis after 96 hours incubation of C2C12 cells with the single agents alone or the combination of vincristine or doxorubicin with the corresponding BH3 mimetics S63845 and A-1331852 for 96 hours. Values indicate mean values ± SEM. All experiments were performed at least three times.*

### Figure 2

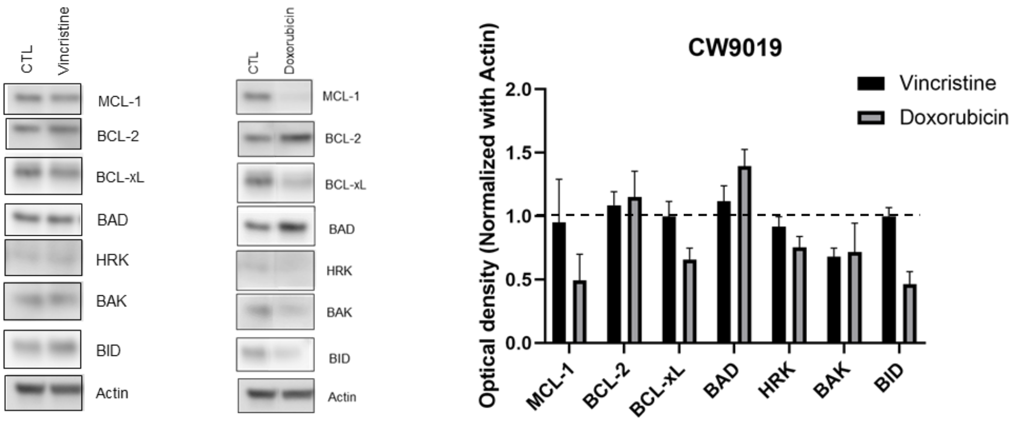


***Supplementary Figure 2: Dynamic BH3 profiling predicts chemotherapy sensitivity in RD and RH4 cell lines.*** *Left panel: images from Western blot analysis from control CW9019 cells and after the treatment with vincristine (left) and doxorubicin (right) for 36 hours. Right panel: Quantification of the optical density of each protein and normalized with actin. Results expressed as fold increase represents the increase in optical density compared to control cells.*

**Figure 3**

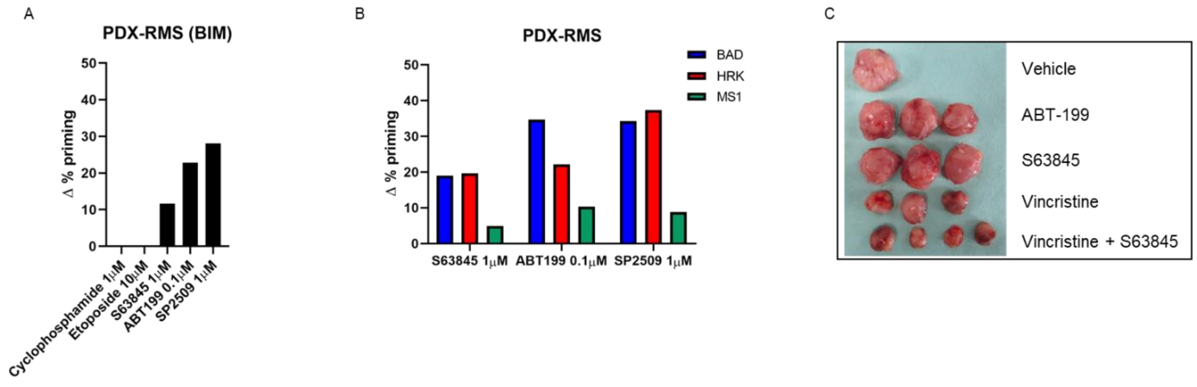


***Supplementary Figure 3: Identification of effective treatments as single agents and combinations with BH3 mimetics in a PDX model of RMS.*** *(A) DBP results of PDX cells from RMS cancer patient showing an increase in Δ% priming after S63845, ABT199 and SP2509 but not after cyclophosphamide or etoposide treatment. Results expressed as Δ% priming represents the increase in priming compared to control cells. (B) DBP results of PDX cells from RMS cancer patient with the sensitizer peptides HRK BH3 peptide and MS1 BH3 peptide showed an increase after all treatments, indicating BCL-xL and MCL-1 adaptation. (C) Image of tumors dissected from PDX mice after 25 days of treatment.*
